# Effectiveness of physical inactivation methods of dengue virus: heat-versus UV-inactivation

**DOI:** 10.1101/427666

**Authors:** Fakhriedzwan Idris, Siti Hanna Muharram, Zainun Zaini, Suwarni Diah

## Abstract

**Introduction:** Complete inactivation of virus is crucial before samples are manipulated outside of biological containment areas or general cleaning. There are several control methods that could decrease the risk from viral contamination on surfaces, which include chemical disinfectants, heating sterilisation, and ultraviolet germicidal irradiation depending on the nature and properties of the materials to be sterilised. To date, there are limited studies reporting the effectiveness of physical inactivation methods of dengue virus. Therefore, this study was designed to evaluate the effectiveness of two physical inactivation methods, which are heat- and ultraviolet-inactivation, against dengue virus.

**Material and methods:** All dengue virus serotypes were subjected to heat treatment at various temperatures and exposed to UV light (wavelength of 250-270 nm) at a distance of approximately 75 cm in a Class II Biosafety cabinet (ESCO) at room temperature. The effectiveness of inactivation methods was tested using viability testing on Vero cells and immunofluorescence assay.

**Results:** Dengue virus can be effectively inactivated by heat treatment at 56°C for at least 30 minutes or at higher temperature. On the other hand, the virus required 45 minutes or longer of ultraviolet light exposure at 75 cm distance from the source to be completely inactivated.

**Conclusion:** The results indicated that DENV can be effectively inactivated using high temperature, i.e. 56°C or above, and UV light irradiation. This result would serve as guidelines in proper decontamination and control of dengue virus in laboratory settings, provided proper conditions are met.

## Introduction

Dengue virus (DENV) is a member of the *Flaviviridae* family with four distinct serotypes, designated as DENV1 to 4. Annually, 50 million people worldwide are estimated to contract dengue virus, and approximately 500,000 to 1,000,000 infections lead to dengue haemorrhagic fever (DHF) or dengue shock syndrome (DSS) with 5 to 30% mortality rates.^1^ With no licensed vaccines as well as specific antiviral treatments available to prevent dengue infection, dengue is considered a major public health problem in the subtropical and tropical regions. Due to this, research in the field is still active and the use of live virus is essential in laboratories. Complete inactivation of virus is crucial before samples are manipulated outside of biological containment areas or general cleaning as well as for research purposes. There are several control methods that could decrease risk from viral infection on surfaces, which include chemical disinfectants, heating sterilisation, and ultraviolet (UV) germicidal irradiation depending on the nature and properties of the materials to be sterilised.^2^ To date, there are limited studies reporting the effectiveness of inactivation methods on DENV. Therefore, this study was designed to evaluate the effectiveness of two physical inactivation methods (heat- and UV-inactivation) against DENV. The outcomes of this study would offer evidences for proper virus inactivation methods and control measures after handling the live viruses.

## Materials and methods

### Cells and virus culture

Clinical isolates used in this study were obtained from Virology Laboratory, Clinical Laboratory Services, Ministry of Health, Brunei Darussalam. DENV isolates were propagated in Vero cells, harvested and stored at −80°C in small aliquots and was used as the source of virus for all experiments.

Vero CCL-81 (African green monkey) were purchased from American Type Culture Collection (ATCC). Vero was grown in MEM (ATCC) containing Earle’s salts with 25 mM HEPES and sodium bicarbonate, supplemented with 10% FBS (SIGMA), 2 mM L-glutamine (SIGMA) and 100 I.U. penicillin-streptomycin ml^−1^ (SIGMA). One-half of the medium was changed every third or fourth day. The cell growth and viability were maintained under constant conditions of 37°C, 5% CO_2_ and a humidified atmosphere in a cell culture incubator.

The viruses were propagated in Vero cells monolayer in a 37°C humidified incubator with 5% CO_2_ and harvested after 7 days inoculation. The virus titres, determined as pfu ml^−1^, were evaluated according to Schaffer et al.^3^ The virus suspension was aliquoted into single-use tube and stored at −80°C until further use.

## Physical inactivation of DENV

### Heat inactivation

DENV serotype 1-4 were subjected to the heat at various temperatures. 150 μl virus stock containing 2.29 × 10^3^ pfu ml^−1^ were transferred into thin-walled 0.2 ml polypropylene tubes and placed in a water bath at different temperatures, including 56 and 70°C for 15, 30, 45, and 60 minutes and additionally, 121°C for 20 minutes in glass bijou bottles using an autoclave (Hirayama). The treated virus was then placed in an ice bath to stop the heat treatment and inoculated to Vero cell monolayer to test the inactivation efficacy.

### UV-inactivation

50μl of DENV serotype 1-4 stocks containing 2.29 × 10^3^ pfu ml^−1^ was spread into a thin layer on 6-well plate (Costar, Corning) and exposed to UV light (wavelength of 250-270 nm) at a distance of approximately 75 cm in a Class II Biosafety cabinet (ESCO) at room temperature. The cover of the 6-well plate was removed when samples were treated with UV light. 450 μl of maintenance medium (Eagle’s Minimum Essential Medium containing 1000 mg glucose L^−1^, 110 mg sodium pyruvate L^−1^ and sodium bicarbonate, supplemented with 2% fetal bovine serum, 2 mM L-glutamine and 100 I.U. penicillin-streptomycin ml^−1^) was added to the well after UV exposure for 15, 30, 45 and 60 minutes respectively. The efficacy of the inactivation process was tested on Vero cell monolayer.

### Virus inactivation testing in Vero cell monolayer

All treated viruses were inoculated into confluent Vero cell monolayer (1.0 × 10^5^ cells ml^−1^ seeding concentration) grown in 6-well plate to determine their infectivity. Each of the treated virus suspension was diluted 1:50 in maintenance medium as well as untreated virus stock as positive control. The monolayer treated with the suspensions were incubated for 4 days at 37°C with 5% CO_2_ and the number of wells developing cytopathic effect (CPE) were recorded.

### Immunofluorescence assay

The treated Vero cell monolayers grown on sterile coverslip were first fixed in ice cold acetone for 5 minutes. The monolayers were then permeabilised in permeabilisation buffer containing 0.2% TritonX-100 and 2% BSA in PBS for 30 minutes. Incubation with rabbit polyclonal anti-DENV 1+2+3+4 antibody (Abcam) was then performed for 1 hour at room temperature. The antibody was diluted 1:100 in PBS containing 10mM glycine, 0.05% Tween20, 0.1% TritonX-100, and 0.1% hydrogen peroxide. This was followed by incubation with the goat anti-rabbit secondary antibody Alexa 488 conjugate (Invitrogen). The monolayer was counterstained with DAPI (Cell Signaling) and mounted with DAKO fluorescence mounting medium (Dako). Three times washing in PBS were done between each step. Imaging was performed using epifluorescence microscope Nikon Eclipse 90i equipped with a Nikon DS-Fi1 camera.

## Results

The properties of DENV were investigated involving the stability of virus when subjected to different heat treatments and UV exposure. In the heat-stability evaluation, all serotypes were subjected to heat treatment at 56°C, 70°C and 121°C either in a wet or pressurised settings. As shown in Table 1, two out of six wells containing Vero cells monolayer of DENV1 and 3, three out of six wells of DENV2, and 4 out of six wells of DENV4 showed CPE under the treatment of 56°C for 15 minutes, indicating the viruses could tolerate these conditions. However, treatments for a longer duration or higher temperature effectively killed the virus, i.e. no viable virus was detected. This observation was confirmed by immunofluorescence assay using DENV1 as the representative serotype. DENV1 particles were undetectable (Figure 1B) compared to the control (Figure 1A). To determine the efficacy of UV treatment in inactivating DENV contaminated surfaces, viruses were spread on 6-well plate and exposed to UV light at varied times at room temperature. The results showed that all serotypes were completely inactivated after 45 minutes UV exposure or longer as shown in Table 2. It should be noted that under 15 minutes UV exposure, three out of six wells were CPE-positive with DENV1, five out of six wells were CPE-positive with DENV2 and 4, and four out of six wells were CPE-positive with DENV3.

**Table 1.**
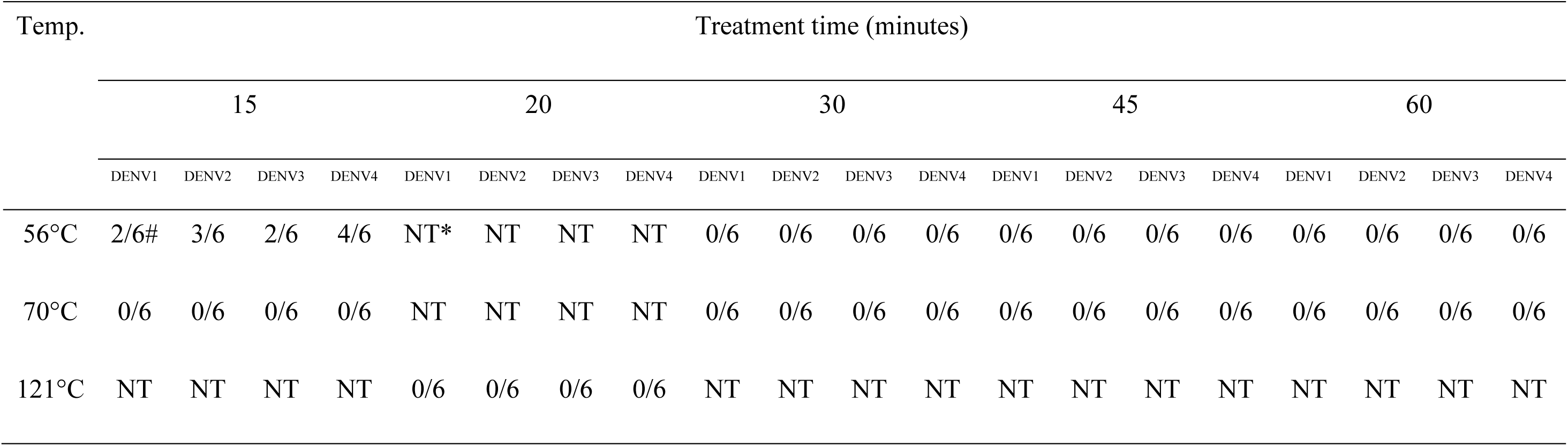
Dengue virus viability under heat treatments.

| Temp. | Treatment time (minutes) |  |  |  |  |  |  |  |  |  |  |  |  |  |  |  |  |  |  |  |
| --- | --- | --- | --- | --- | --- | --- | --- | --- | --- | --- | --- | --- | --- | --- | --- | --- | --- | --- | --- | --- |
|  | 15 |  |  |  | 20 |  |  |  | 30 |  |  |  | 45 |  |  |  | 60 |  |  |  |
|  | DENV1 | DENV2 | DENV3 | DENV4 | DENV1 | DENV2 | DENV3 | DENV4 | DENV1 | DENV2 | DENV3 | DENV4 | DENV1 | DENV2 | DENV3 | DENV4 | DENV1 | DENV2 | DENV3 | DENV4 |
| 56°C | 2/6# | 3/6 | 2/6 | 4/6 | NT* | NT | NT | NT | 0/6 | 0/6 | 0/6 | 0/6 | 0/6 | 0/6 | 0/6 | 0/6 | 0/6 | 0/6 | 0/6 | 0/6 |
| 70°C | 0/6 | 0/6 | 0/6 | 0/6 | NT | NT | NT | NT | 0/6 | 0/6 | 0/6 | 0/6 | 0/6 | 0/6 | 0/6 | 0/6 | 0/6 | 0/6 | 0/6 | 0/6 |
| 121°C | NT | NT | NT | NT | 0/6 | 0/6 | 0/6 | 0/6 | NT | NT | NT | NT | NT | NT | NT | NT | NT | NT | NT | NT |

**Figure 1.**
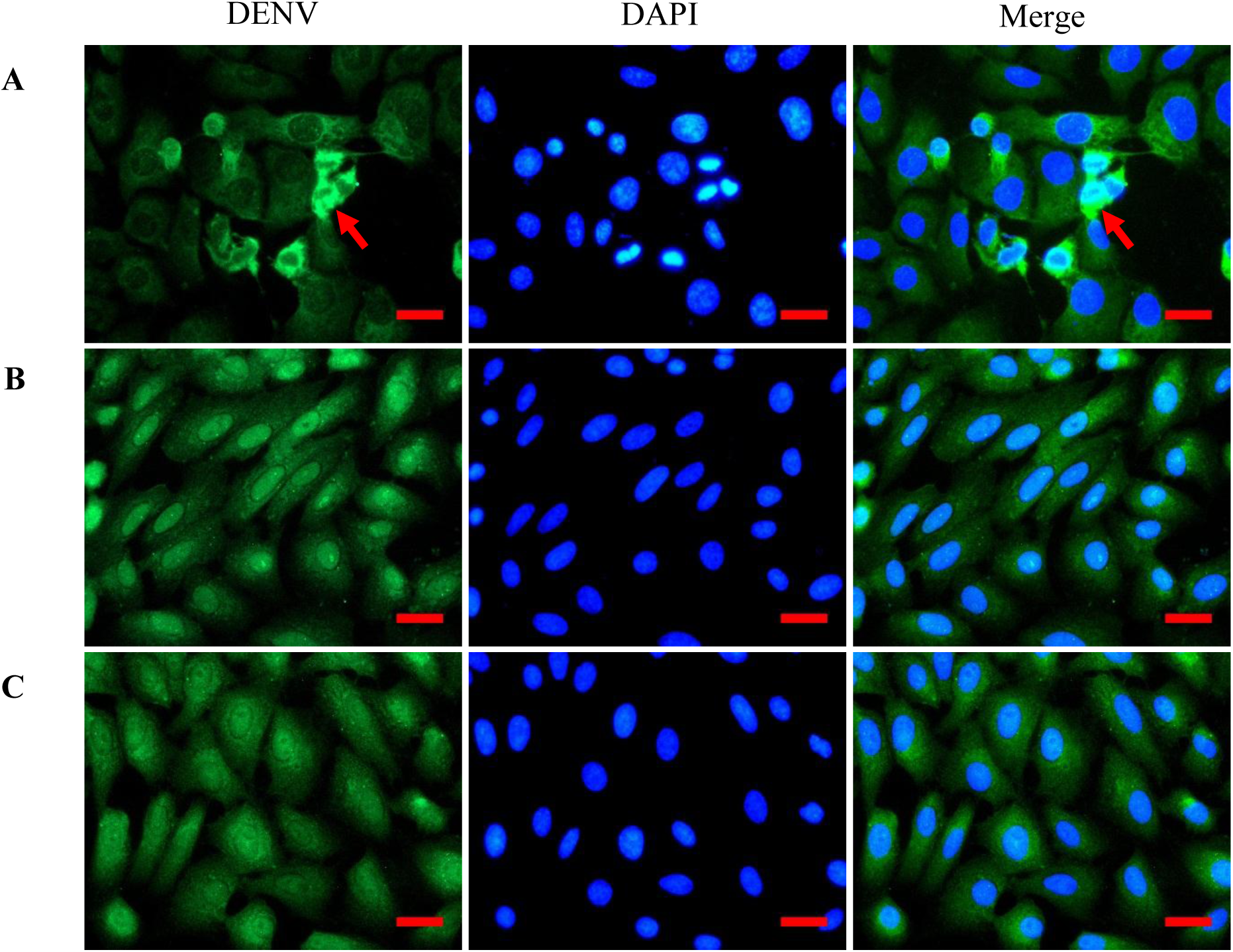
Detection of DENV1 in Vero cells to evaluate the efficacy of different physical treatments (X40 magnification). DENV was targeted using anti-DENV 1+2+3+4 polyclonal IgG antibodies (Abcam) and counter-stained using AlexaFluor 488 conjugated secondary antibody (Abcam) (red arrows). Nuclei were stained using DAPI (Cell Signalling). A) control; Vero cells were inoculated with untreated DENV1 suspension, B) Heat-inactivated DENV1 suspension at 121°C for 20 minutes, C) 60 minutes UV-inactivated DENV1 suspension. Image merging was done using FIJI/ImageJ software. Scale = 50μm.

**Table 2.** Dengue virus viability under UV irradiation treatment.

| Serotype | DENV1 |  |  |  | DENV2 |  |  |  | DENV3 |  |  |  | DENV4 |  |  |  |
| --- | --- | --- | --- | --- | --- | --- | --- | --- | --- | --- | --- | --- | --- | --- | --- | --- |
| Exposure time (minutes) | 15 | 30 | 45 | 60 | 15 | 30 | 45 | 60 | 15 | 30 | 45 | 60 | 15 | 30 | 45 | 60 |
| No. of positive wells | 3/6* | 1/6 | 0/6 | 0/6 | 5/6 | 4/6 | 0/6 | 0/6 | 4/6 | 2/6 | 0/6 | 0/6 | 5/6 | 1/6 | 0/6 | 0/6 |

Note: \*Infectivity to Vero cell monolayers in wells = (Infected wells)/(total wells inoculated; n = 6) per treatment.

## Discussion

The aim of this study was to validate the two commonly used physical inactivation methods against DENV for protection or the decontamination methods at laboratory or field conditions. The results indicated that all serotypes can be effectively inactivated by temperature treatment at 56°C for 30 minutes and at higher temperatures. But, it should be noted that DENV still could survive after treatment at 56°C for 15 minutes. To date, this study is the first to report the heat treatment and stability of DENV of clinical isolates. However, our data are in agreement with other studies done on other members of *Flaviviridae* which may reflect the similarities among the members. For example, TBEV was completely deactivated at 72°C for 15s which simulates the pasteurisation method.^4^ Alkhumra hemorrhagic fever virus infectivity was lost completely when heat-treated at 60°C for 3 minutes and 56°C for 30 minutes.^5^ Comparable results were observed with yellow fever virus which was completely inactivated by heating at 56°C for 30 minutes and 60°C for 5 minutes.^6^ Similar outcomes have also been reported with St. Louis encephalitis virus, WNV, and hepatitis C virus.^7,8^ The loss of infectivity of the virus was owed to the structural changes in viral proteins induced by heat and viral RNA degradation.^9-12^

The present study also showed that UV treatment of DENV serotypes was effective when exposed for 45 minutes or longer within 75 cm distance. Nevertheless, the virus was able to survive if exposed to UV for shorter than 30 minutes. The result does not only indicate the UV tolerance of DENV but also suggested that sufficient UV exposure time is necessary to avoid cross-contamination on lab-works or general surface sterilisation. UV irradiation, typically at a wavelength of 254nm, is reported to target nucleic acids impairing the replication capability while leaving proteins mostly preserved.^9,13^

## Conclusion

The results indicated that DENV can be effectively inactivated using high temperature, i.e. 56°C or above, and UV light irradiation. This result would serve as guidelines in proper decontamination and control of DENV in laboratory settings as well as for research purposes such as vaccine development, provided proper conditions are met.

## Acknowledgements

The authors would like to thank Virology Laboratory, Clinical Laboratory Services, Laboratory Services, Ministry of Health, Brunei Darussalam for allowing the use of dengue virus stock for this study. Idris F is a recipient of the Graduate Research Scholarships (GRS), Universiti Brunei Darussalam.

## Compliance with Ethics Guidelines

This article does not contain any studies with human or animal subjects performed by any of the authors

## Author contributions

FI, SHM, ZZ and SD conceived the experiments. FI performed the experiments. FI, SHM, ZZ and SD analysed the results, and FI wrote the first version of the manuscript. SHM, ZZ and SD checked and finalised the manuscript.

